# mHMG-DTI: a drug-target interaction prediction framework combining modified Hierarchical Molecular Graphs and improved Convolutional Block Attention Module

**DOI:** 10.1101/2024.12.21.628851

**Authors:** Zerui Yang, Yinqiao Li, Yudai Matsuda, Linqi Song

## Abstract

Drug-target interactions (DTIs) are fundamental to understanding the therapeutic mechanisms of drugs, yet accurately predicting these interactions remains a significant challenge in drug discovery. Current computational approaches often fail to capture essential molecular motifs and spatial information of proteins, limiting their effectiveness, particularly when encountering proteins or compounds absent in the training datasets. To address these limitations, we propose mHMG-DTI, a novel framework that leverages an improved Convolutional Block Attention Module (iCBAM) for enhanced protein feature extraction and modified Hierarchical Molecular Graphs (mHMGs) for comprehensive molecular encoding. This hierarchical approach not only captures detailed local structures and broader connectivity patterns but also incorporates guiding knowledge to improve feature representation. Across a total of 16 experimental evaluations on four benchmark datasets spanning both classification and regression tasks, mHMG-DTI surpasses existing baseline models in 11 cases. These results highlight the potential of mHMG-DTI to enhance DTI prediction accuracy, thereby accelerating the drug discovery process and providing valuable insights into drug resistance and side effect mechanisms.

## 1 Introduction

Drug-target interaction (DTI) studies are critical for elucidating the mechanisms through which drugs produce their therapeutic effects. These interactions can manifest in various forms, such as direct drug binding to target proteins (Zhao et al., 2022), modulation of protein functionality (Wu et al., 2020), or changes in protein expression levels (Li et al., 2021). The precise identification and characterization of DTIs are essential for drug discovery, development, and optimization, as well as for understanding potential side effects and the development of drug resistance.

A range of experimental techniques, including biochemical assays (Bates et al., 2024), X-ray crystallography (Kermani, 2020), and nuclear magnetic resonance (NMR) spectroscopy (Reif et al., 2021), are commonly employed to detect and analyze DTIs. However, these experimental methods are often labor-intensive, time-consuming, and resource-demanding. In response to these limitations, deep learning (DL) has gained prominence as a powerful tool for DTI prediction. DL algorithms, particularly neural networks, are adept at capturing complex, non-linear patterns within large datasets, making them especially suited to the intricacies of DTI prediction. Consequently, several computational approaches have been developed to streamline and accelerate this process, offering significant advantages in terms of efficiency and scalability.

The study by Lee et al. (2019) introduces DeepConv-DTI, which utilizes convolutional layers to extract protein features and the extended-connectivity fingerprint (ECFP) (Rogers and Hahn, 2010) for molecular representation before integration into fully connected layers. This method marks a shift from the traditional use of fingerprints or descriptors, as convolutional neural networks (CNNs) have proven effective in extracting molecular information by treating molecules as plain text. In another advancement, HyperAttentionDTI (Zhao et al., 2021) employs a sequencebased deep learning framework with an attention mechanism to enhance predictions of DTIs. This model combines CNNs with an attention mechanism to discern complex interactions at both atomic and amino acid levels. Following the introduction of the Transformer architecture by Vaswani et al. (2017), Shin et al. (2019) proposed MT-DTI, and Huang et al. (2020) introduced MolTrans. Both models exploit the self-attention mechanism to effectively encode molecular structures. A primary limitation of these computational approaches in drug discovery is their inadequate effectiveness in providing comprehensive molecular information. Owing to the intrinsic topological structure of molecules, where each chemical bond is conceptualized as an edge and each atom as a node, Nguyen et al. (2020) exploited Graph Neural Networks (GNNs) to elucidate molecular representations. The seminal work presented in the GraphDTA study introduces a pioneering method for predicting drug-target binding affinity using GNNs.

Contemporary research underscores the critical importance of molecule and protein representations in predictive modeling, particularly within DTI studies. Three principal strategies have emerged for encoding molecules in DTI interaction predictions: one-dimensional convolutional neural networks (1D-CNNs) (Öztürk et al., 2018), self-attention mechanisms (Chen et al., 2020), and GNNs (Yazdani-Jahromi et al., 2022; Bai et al., 2023). However, these methods frequently fall short of capturing local molecular features, such as motifs, rings, and functional groups, which are crucial for creating informative molecular representations. As a result, existing molecular encodings may limit the performance of predictive models. In addition, both 1D-CNNs and GNNs face difficulties in identifying long-range atomic interactions, which are typically confined to nearestneighbor exchanges during the message-passing process (Wu et al., 2022). In addition, many approaches also underutilize prior chemical knowledge, further hindering performance. For example, the DeepConv-DTI model relies on Morgan fingerprints for molecular encoding, followed by a fully connected layer, thus neglecting the molecular topology. On the other hand, Perceiver CPI (Nguyen et al., 2022) integrates molecular descriptors and graph structures via cross-attention, offering a more holistic approach.

In the field of protein representation, there is ongoing debate regarding the use of sequence-based versus graph-based models. Sequence-based models (Zhao et al., 2021; Bai et al., 2023), although efficient, often fail to capture essential structural features, while graph-based models (Jiang et al., 2020; Jiang et al., 2022), though more informative, significantly increase computational complexity without consistently improving performance (Zhang et al., 2023). A notable attempt to bridge this gap is the saCNN model (Wang et al., 2021), which integrates the Convolutional Block Attention Module (CBAM) to capture spatial dependencies within proteins. However, in their implementation, the channel attention component of CBAM was removed due to its negative impact on model performance. In contrast, our study demonstrates that the inclusion of channel attention is effective for learning protein input features and enhancing model robustness.

To address these challenges, we propose mHMG-DTI (Drug-Target Interaction prediction with modified Hierarchical Molecular Graphs), a novel framework inspired by the Hierarchical Molecular Graphs (HMGs) introduced by Zang et al. (2023) for molecular encoding. HMGs are specifically designed to capture substructural motifs through a hierarchical representation. While a similar approach has been explored by Liu et al. (2024), their work formulates DTI prediction solely as a classification task, lacking comprehensive analysis. Additionally, their method largely replicates the original framework, initializing the global level node from scratch, which, as our experiments demonstrate, can be further optimized.

Building upon the foundational HMG framework, we introduce a modified representation by incorporating guiding knowledge into the graph structure, thereby improving the extraction of critical molecular information. For protein encoding, our model integrates recent advances in attention mechanisms (Chen et al., 2016; Jia et al., 2023) and employs an improved Convolutional Block Attention Module (iCBAM) to optimize feature extraction across both channel and spatial dimensions. The information exchange between molecular and protein representations is facilitated through two cross-attention modules, enabling enhanced interaction learning. Furthermore, we show that the incorporation of large biological models significantly boosts the overall predictive performance of the proposed framework.

The key contributions of this work are as follows:

1. To the best of our knowledge, our model represents the first application of Hierarchical Molecular Graphs for molecular representation in DTI prediction of both classification and regression task, effectively capturing molecular motifs.
2. We extend the original HMG framework by embedding molecular descriptors as auxiliary information, facilitating seamless integration with graph neural networks (GNNs) without the need for additional network architectures.
3. We introduce a novel application of iCBAM for protein representation in the context of DTI prediction, achieving improved feature extraction and model performance.

Our experimental results demonstrate that mHMG-DTI outperforms baseline models across multiple benchmark datasets, setting a new standard in DTI prediction. This work provides a comprehensive and accurate framework for addressing limitations in current methodologies, offering promising implications for drug discovery.

## 2 Materials and method

### 2.1 Datasets

In this study, we formulate DTI prediction as both a classification and a regression task. For the regression task, we utilize the KIBA (Tang et al., 2014) and Metz (Metz et al., 2011) datasets, while for the classification task, we employ BindingDB (Gilson et al., 2015) and DrugBank (Wishart et al., 2005).

The KIBA dataset (Kinase Inhibitor BioActivity) integrates data from large-scale kinase inhibitor bioactivity profiling studies, which measure various bioactivity parameters, including IC_50_, *K*_*i*_ (inhibition constant), and *K*_*d*_ (dissociation constant). KIBA consolidates and normalizes these diverse measurements into a single bioactivity score, offering a unified and comprehensive metric for describing the interactions between kinase inhibitors and their targets, thereby enhancing traditional DTI prediction methods. The Metz dataset compiles molecular data for chemical and biological tasks, selecting molecules from databases and measuring properties such as binding activity and toxicity as ground truth labels. The BindingDB dataset derives data primarily from scientific articles and US patents, detailing at least one protein target, a small molecule compound, and a quantitative measure of their interaction (e.g., IC_50_, *K*_*i*_, or *K*_*d*_). Following the procedure described by Gao et al. (2018), the dataset was transformed into a binary classification format. The DrugBank dataset was adapted from Zhao et al. (2021), with inorganic compounds, very small molecules, and molecules with SMILES strings unrecognized by RDKit removed during preprocessing. The detailed information on the datasets can be found in Table 1, Figure 1, and Supplementary Figures S1 and S2.

**Table 1.** Statistic of the datasets of the classification task.

| Dataset | Positive | Negative | Total |
| --- | --- | --- | --- |
| BindingDB | 33772 | 27486 | 61258 |
| DrugBank | 17511 | 17511 | 35022 |

**Figure 1.**
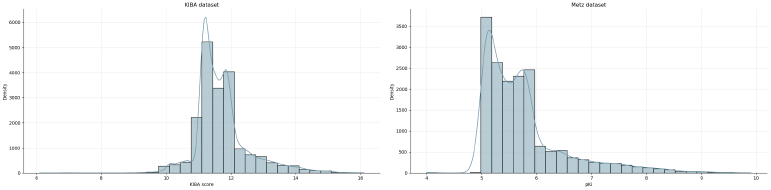
Statistics of the datasets of the regression task.

### 2.2 Model architecture

The proposed model architecture (Figure 2) integrates molecular and protein representations by combining graph neural networks (GNNs), convolutional attention mechanisms, and crossattention blocks, while leveraging pre-trained large model (PLM) embeddings for both molecules and proteins. The GNN module, which processes molecular data, includes two input embedding layers to encode atom types and degrees, as well as two additional embedding layers for bond types and bond directions. Molecular data is propagated through a series of Graph Convolutional Networks (GCN) to capture the structural information of the molecules. To enhance the expressiveness of the graph, self-loops are incorporated, ensuring that each node is connected to itself. Additionally, edge attributes are included to enrich the molecular graph, which is subsequently utilized to update the molecular embeddings.

**Figure 2.**
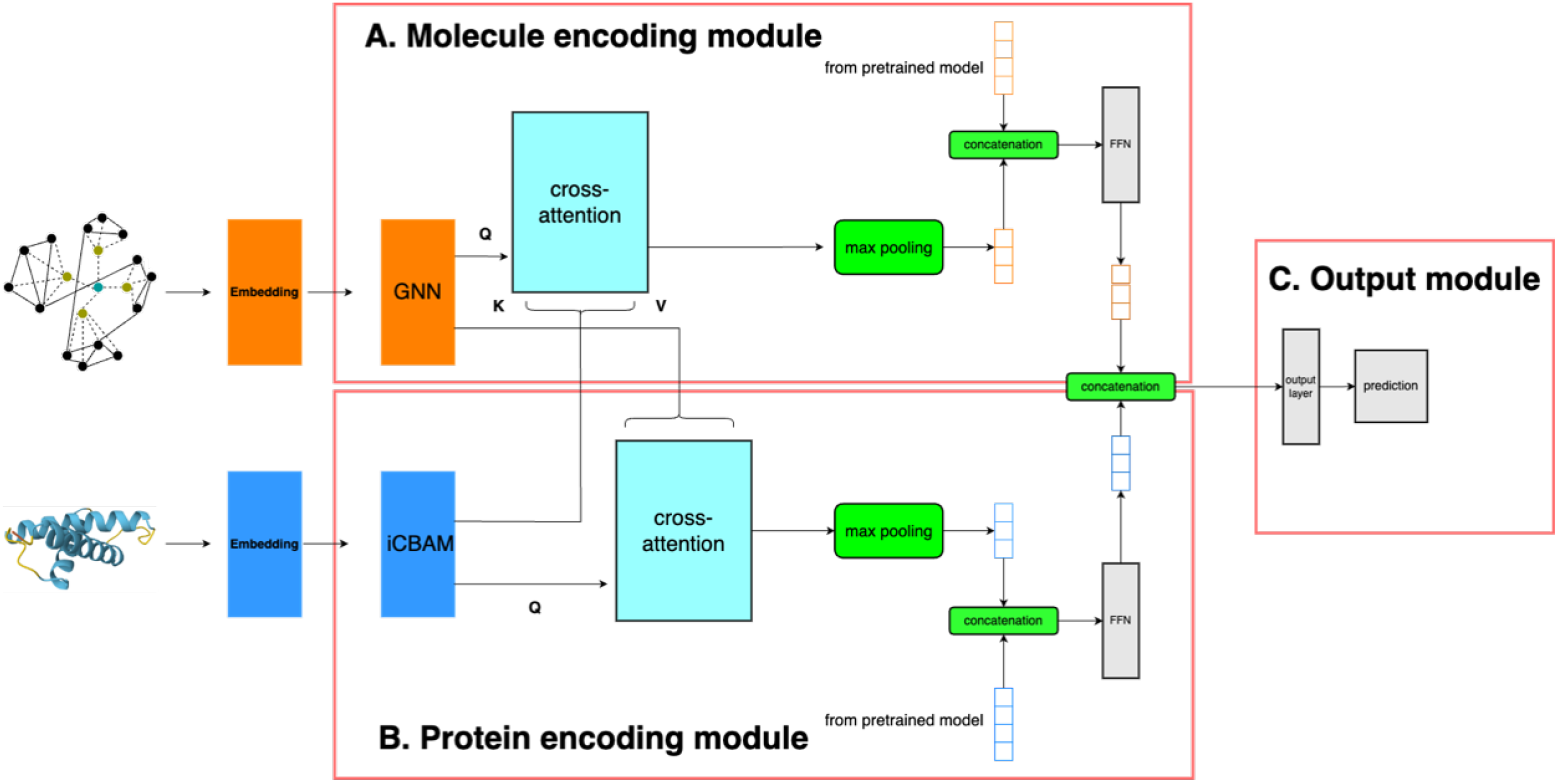
Structure of HMG-DTI.

For processing 2D protein structures, the model employs a ResNet (He et al., 2015) layer comprising several convolutional layers with skip connections, which preserve learned information across layers. Following the convolutional layers, Spatial Attention and Channel Attention mechanisms are applied in parallel to selectively emphasize significant spatial and channel-wise features within the protein data. These attention modules dynamically adjust the model’s focus, thereby improving its capacity to capture critical features.

To facilitate the exchange of information between molecular and protein features, a multihead cross-attention mechanism is employed. The cross-attention is computed bidirectionally: molecules are treated as queries (Q) and proteins as keys (K) and values (V ), and vice versa. Residual connections are added to enhance information flow and stability during learning.

After processing through their respective modules, the molecular and protein representations undergo max-pooling and are concatenated with their corresponding pre-trained LLM encodings. These concatenated representations are then passed through linear projection layers for dimensionality reduction. Finally, the molecular and protein representations are combined and fed into the output module to generate the final predictions.

### 2.3 Hierarchical Molecular Graph

The previous methods (Sun et al., 2021) (Wang et al., 2022) for learning molecular representations focused primarily on the atomic level, with each atom depicted as a node and each chemical bond as a link. These methods have a significant limitation; they overlook the motif structure of the molecule, which contains crucial information about the molecule. Furthermore, these approaches are unable to represent the molecule holistically, capturing only localized information instead.

In this study, we employ a Hierarchical Molecular Graph (Zang et al., 2023) to encode the molecule more comprehensively. To be detailed, the graph of the molecule is constructed as *G* = (V_*atoms*_, *E*_*a*_) according to its chemical structure, where *V*_*atoms*_ denotes the atoms, and *E*_*a*_ denotes the chemical bonds. Using the BRICS algorithm (Liu et al., 2017), the molecule is first broken down into various motifs, while the remaining skeleton is further decomposed into individual rings. Both motifs and rings are then represented by augmented motif nodes 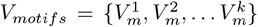, which connect to their respective atom nodes *V*_*atoms*_ forming a new set of bonds *E*_*m*_. Ultimately, an augmented graph node *V*_*graph*_ is created and linked to all motif nodes *V*_*motifs*_, forming the bonds *E*_*g*_. All augmented nodes and bonds are integrated into the original graph to form the augmented graph *G*_*aug*_ = (V, *E*), where *V* = {V_*atoms*_, *V*_*motifs*_, *V*_*graph*_} and *E* = {E_*a*_, *E*_*m*_, *E*_*g*_}. Each *V*_*atoms*_ has two properties, atomic number and degree, while each Ea also has two properties, bond type and a binary value for if it is in a ring. All augmented nodes and edges are assigned the corresponding pseudo values, respectively.

Based on the original algorithm, we make one modification by substituting the value of graph node *V*_*graph*_ (after embedding layer) with Morgan fingerprint as the guiding knowledge instead of learning from scratch. This hierarchical approach allows for a richer and more informative representation of molecules, capturing both the detailed local structures and the broader connectivity patterns that are critical for understanding molecular properties and behaviors. In addition, it incorporates the molecular fingerprints, which contain rich chemical information without additional computation, such as cross-attention (Nguyen et al., 2022), enhancing the computational efficiency of the model.

### 2.4 ResNet

The core innovation of ResNet lies in its use of skip connections, also known as residual connections, which allow information to bypass one or more layers. This design facilitates the training of very deep networks by providing a mechanism through which gradients can flow more easily back through the network during backpropagation. The basic building block of a ResNet is the residual block, which typically consists of two parts:

1. Identity Mapping (Skip Connection): An identity mapping is introduced where the output of a layer is added to the output of a previous layer. This can be represented mathematically as *H*(*x*) = *F* (*x*) + *x*, where *H*(*x*) is the desired underlying mapping to be learned, and *F* (*x*) is the learned residual mapping.
2. Convolutional Layers: These layers perform the standard convolution operations on the input data. They learn the necessary features that, when combined with the identity mapping, form the final output of the block.

ResNet has been intensively used for the recognition of the pattern of biological molecules. However, in most cases, 1D convolutional layers were applied by treating the biological molecules as plain text without any changes in the channel number. In our model, the ResNet layer is constructed by stacking three of these residual blocks sequentially with 2D convolutional layers through which the channel number of protein input data expands from one to sixteen to capture more abstract features, which we consider as augmented spatial features.

### 2.5 Channel attention module

The Channel Attention (CA) module is a pivotal component in modern convolutional neural networks (CNNs), designed to improve feature representation by emphasizing informative channels and suppressing less useful ones, thereby enhancing the discriminative power of the network. Given a feature map *X* ∈ *R*^*C∗H∗W*^ , where C denotes the number of channels and H and W represent the height and width, the CA module first aggregates spatial information through global average pooling (GAP) and global max pooling (GMP), yielding a channel descriptor *z* ∈ *R*^*C*^; this descriptor is then passed through a series of fully connected (FC) layers serving as a bottleneck, which reduce the dimensionality to C/r and subsequently expand it back to C, with a non-linear activation function (e.g., ReLU) introduced between these layers. The output is normalized using an activation function like Sigmoid to ensure values lie within the range [0, 1], generating a channel attention map *M*_*c*_ ∈ *R*^*c*^; finally, this map is multiplied elementwise with the original feature map X to produce the weighted feature map X’.

### 2.6 Spatial attention module

The Spatial Attention (SA) module is a critical architectural component designed to enhance the efficacy of feature extraction by emphasizing salient regions within an input feature map. Given a protein feature map *X* ∈ *R*^*C∗H∗W*^ , where C denotes the number of features, and H and W represent the sequence length or structural dimensions, the SA module initiates by processing the feature map to extract spatial or positional information. This is typically achieved through a convolution operation, followed by a normalization step such as batch normalization to stabilize the learning process. The processed feature map is then subjected to spatial aggregation, commonly through global average pooling (GAP) and global max pooling (GMP) along the spatial dimensions, summarizing the spatial context and resulting in two sets of feature statistics. These statistics are concatenated along the channel axis and passed through a convolutional layer, often with a kernel size of 7×7, to generate a spatial attention map *M*_*s*_ ∈ *R*^*H∗W*^ . To ensure the attention map values are bounded between 0 and 1, a sigmoid activation function is applied, producing a normalized spatial attention map. The spatial attention map Ms is subsequently used to recalibrate the original feature map *X* via element-wise multiplication, effectively amplifying the features in the highlighted regions while attenuating those in less important areas.

### 2.7 Pretrained large biological model

The ESM-2 model is a cutting-edge large language model designed for protein structure prediction, trained on approximately 65 million unique protein sequences from the UniRef50 database. Using the Masked Language Modeling (MLM) objective, it predicts masked amino acids, enabling the model to capture evolutionary patterns and dependencies within sequences. Ranging from 80 million to 15 billion parameters, ESM-2 utilizes a BERT-based Transformer architecture with enhancements like Rotary Position Embeddings (RoPE) to handle long sequences effectively. Notably, ESM-2 delivers high-resolution predictions at speeds up to 60 times faster than prior stateof-the-art methods, while maintaining accuracy comparable to existing techniques. It supports large-scale predictions, such as for 617 million proteins in the MGnify90 database, and can predict novel protein structures that may provide new insights into function. Furthermore, ESM-2 generates high-confidence predictions that are comparable to experimentally determined structures in terms of functional and biochemical site analysis. These advancements make ESM-2 a powerful tool for large-scale protein sequence embedding.

ChemBERTa is a transformer-based model designed for chemical and molecular representation. It is trained using SMILES strings in a selfsupervised fashion, employing masked language modeling (MLM) to predict masked tokens and learn contextual relationships within molecular structures. The model is exposed to large-scale chemical datasets, allowing it to capture complex molecular dependencies. One of its key advantages is its ability to generalize well across a range of downstream tasks, such as molecular property prediction, with minimal labeled data. Chem-BERTa’s architecture enables it to handle long-range dependencies, making it a robust tool for chemical informatics and drug discovery applications.

### 2.8 Cross-attention

Cross-attention is a mechanism in neural networks, particularly in transformer models, that allows one sequence (query) to focus on relevant parts of another sequence (key and value). The core idea is to compute attention weights between the query sequence and the key-value pair, which then help in producing contextually aware representations. The mechanism is mathematically described using the attention function, typically expressed as:

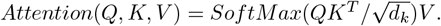

Definitions of terms:

*Q* (Query): A matrix representing the input sequence to which we are applying attention. This matrix is generated by a linear transformation of the input features.

*K* (Key): A matrix representing the elements from which attention is being computed. This matrix is similarly obtained by applying a linear transformation to the input features.

*V* (Value): A matrix whose elements correspond to the key elements, and whose weighted combination produces the final output of the attention mechanism.

*d*_*k*_: The dimensionality of the key vectors. The division by dk is a scaling factor to ensure the dot product does not grow too large as the dimensionality increases. The term *QK*^*T*^ computes the dot product between each query and all keys, producing an alignment score for each key with respect to the query. These alignment scores are then normalized using the softmax function, which transforms the scores into attention weights that sum to one. The output of the attention mechanism is a weighted sum of the values *V* , where the weights are derived from the softmax-normalized alignment scores. The result is a context-aware representation that highlights the most relevant parts of the input based on the query. Mathematically, cross-attention can be extended to multiple heads (MHA), where different subsets of the queries, keys, and values are processed in parallel by separate attention heads:

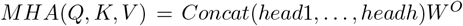

Where each head is defined as:

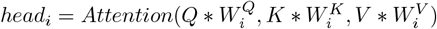

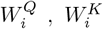 and 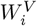 are learnable projection matrices for the query, key, and value, and *W*^*O*^ is a projection matrix for the output of the concatenated heads.

## 3 Result

### 3.1 Experimental Setup

We selected several baseline models originally designed for classification or regression tasks. The output layers of these models were modified to fit the specific task requirements, while the remaining architecture was left unchanged.

A five-fold cross-validation procedure was employed, using an 80% training and 20% test data split. In line with prior research, we defined four distinct experimental settings for partitioning the training and test datasets:

E1: All proteins and drugs in the test set are also present in the training set.

E2: All proteins in the test set are included in the training set, but none of the test set drugs are in the training set.

E3: All drugs in the test set are part of the training set, but no proteins in the test set are present in the training set.

E4: Neither proteins nor drugs in the test set appear in the training set.

### 3.2 Evaluation Metrics

Our tasks involve both classification and regression objectives. For the classification task, we employed several evaluation metrics, including accuracy (ACC), precision (PRE), recall, F1-score, Matthews correlation coefficient (MCC), the area under the receiver operating characteristic (ROC) curve, and the area under the precision-recall (PR) curve. For the regression task, the mean squared error (MSE) and the concordance index (CI) were used to assess performance. All experiments were replicated five times, and the average values were reported to ensure robustness. The comparative performance of our model and baseline models across different scenarios is summarized in Table 1. Overall, our model achieved the best performance in BindingDB dataset under E1-3, in DrugDB dataset under E1-4, in KIBA dataset under E1, E2, E4, in the Metz dataset under E1, E4.

Below, we provide a detailed summary and comparison of our model with other state-of-the-art models, including GraphDTA (Nguyen et al., 2020), DeepConv-DTI (Lee et al., 2019), HyperAttentionDTI (Zhao et al., 2021), and Perceiver CPI (Nguyen et al., 2022). Notably, many of the results obtained in our study closely align with those reported in the original works (e.g., HyperAttentionDTI on the DrugBank dataset under E2 and E3), while some even surpass the original performance (e.g., Perceiver CPI on the Metz dataset under E4).

### 3.3 Experimental Setting Summaries and Model Comparisons

In E1, all proteins and drugs in the test set are also present in the training set. Under these conditions, our model consistently achieved the highest scores across all datasets. For instance, in the BindingDB dataset, our model attained an accuracy of 0.942, surpassing GraphDTA (0.833) and DeepConv-DTI (0.859), while closely matching HyperAttentionDTI (0.921). This indicates that our model can effectively leverage previously seen protein-drug interactions, demonstrating high F1, AUC, and AUPR scores across various datasets.

In E2, all proteins in the test set are included in the training set, but none of the test set drugs are in the training set. Despite this challenge, our model maintains robust performance, particularly in precision and F1 score. For BindingDB E2, our model achieves an accuracy of 0.947, outperforming Perceiver CPI (0.935) and HyperAttentionDTI (0.924). This suggests that our model can generalize well to new drugs when protein information is available, a strength not equally shared by the baseline models, as evidenced by their lower recall values, particularly for GraphDTA (0.800).

In E3, all drugs in the test set are part of the training set, but no proteins in the test set are present in the training set. Here, the model shows a moderate decrease in performance but still maintains higher accuracy than the baselines in many cases. In DrugBank E3, our model’s accuracy of 0.744 surpasses all other baseline models, indicating that our model can generalize well to unseen proteins when drug data is already familiar.

In E4, neither proteins nor drugs in the test set appear in the training set. This is the most challenging setting, yet our model still outperforms other models in certain metrics. For example, in KIBA E4, our model achieves an MSE of 0.551 and CI of 0.634, slightly outperforming DeepConv-DTI (MSE: 0.553, CI: 0.626) and Perceiver CPI (MSE: 0.554, CI: 0.612). HyperAttentionDTI shows comparable results, although it suffers from lower MCC and recall metrics in this difficult setting.

Overall Performance Our model demonstrates robust performance across various dataset scenarios (Figure 3). It outperforms other baseline models in all instances of E1 and in most cases of E2 and E4. We attribute this superior performance to our mHMG molecular representation method, which not only provides information about molecular motifs but also incorporates guiding knowledge that facilitates the model’s learning of molecular information, even in the presence of previously unseen molecules. A similar approach, such as Perceive CPI, employs a combination of molecular graphs and Morgan fingerprints to represent molecules, achieving commendable results in E2, albeit slightly inferior to our model. In E3, our model exhibits a slight decline in performance; however, it still surpasses other models on the DrugBank and BindingDB dataset and demonstrates competitive performance on other datasets. These results (Supplementary Table S1-S16) highlight the adaptability and robustness of our model.

**Figure 3.**
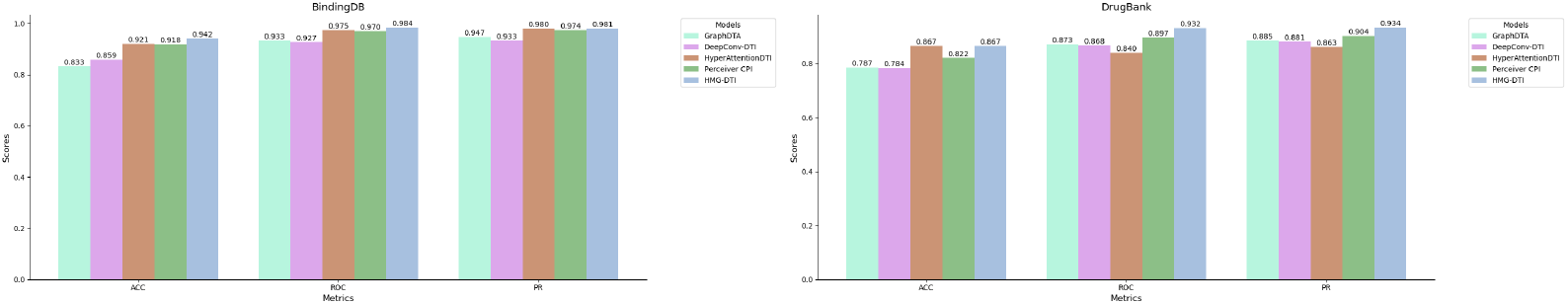
Performance comparison between mHMG-DTI and baseline models under E1.

### 3.4 Ablation studies

To evaluate the contributions of mHMG, iCBAM, and the large model-encoded (LME) molecular and protein representations, we conducted a series of ablation studies (Table 2, Table 3, Table 4). These experiments were structured as follows: (1) comparing the performance of mHMG against standard molecular graph encoding (MG) and the original HMG framework, (2) assessing the effectiveness of iCBAM relative to spatial attention mechanisms, (3) analyzing the impact of excluding LME molecules, and (4) analyzing the impact of excluding LME proteins. These ablation studies comprehensively evaluate the contributions of each component to the overall performance of the proposed framework.

**Table 2.** Ablation studies under E1 where both proteins and molecules in the test set are present in the training set.

| Models | Bindingdb | DrugBank | KIBA | Metz |
| --- | --- | --- | --- | --- |
| mHMG-DTI | 0.942 | 0.867 | 0.361 | 0.186 |
| HMG | 0.936 | 0.865 | 0.362 | 0.190 |
| MG | 0.940 | 0.861 | 0.362 | 0.204 |
| Spatial attention | 0.936 | 0.864 | 0.373 | 0.192 |
| *w/o*_LME_M | 0.931 | 0.858 | 0.410 | 0.234 |
| *w/o*_LME_P | 0.924 | 0.850 | 0.395 | 0.228 |

**Table 3.**
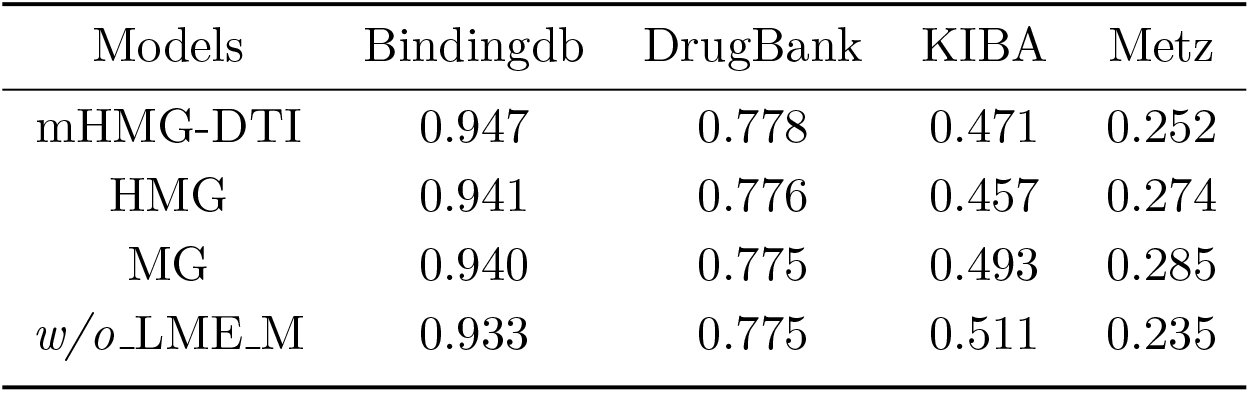
Ablation studies under E2 (with seen proteins and novel molecules).

**Table 4.** Ablation studies under E3 (with novel proteins and seen molecules).

| Models | Bindingdb | DrugBank | KIBA | Metz |
| --- | --- | --- | --- | --- |
| mHMG-DTI | 0.897 | 0.744 | 0.410 | 0.452 |
| Spatial attention | 0.889 | 0.741 | 0.423 | 0.443 |
| *w/o*_LME_P | 0.877 | 0.724 | 0.411 | 0.384 |

Since mHMG, HMG, MG, and LME molecules primarily affect the extraction of molecular information, their ablation experiments were conducted exclusively under E1 and E2. Similarly, the ablation experiments for iCBAM and LME proteins were performed under E1 and E3. The results demonstrated that our model outperformed the control conditions in most cases, with exceptions in the KIBA dataset under E2 and the Metz dataset under E2 and E3. Notably, LME proteins and molecules were more effective in E1, where both proteins and molecules had been observed during training. Furthermore, LME molecules and proteins contributed more significantly to the classification task than to the regression task, with the contribution of LME proteins being substantially larger than that of LME molecules.

The contribution of mHMG was significantly greater than that of iCBAM, which is expected given that mHMG integrates pre-calculated structural information and guiding knowledge, while iCBAM does not provide additional protein-specific information.

Compared to MG, HMG generally exhibited better performance except for the BindingDB dataset under E1. In contrast to LME molecules and proteins, the HMG and mHMG methods demonstrated a higher contribution in regression tasks, which require finer granularity, further highlighting their effectiveness in handling more complex tasks.

## 4 Analysis

We visualized the kernels of the spatial attention convolutional layer and calculated the variance within each kernel (Figure 4, Supplementary Figures 3–5), revealing intriguing patterns. Specifically, the kernels associated with mean pooling channels exhibit lower variance in their activations compared to those associated with max pooling channels. This is likely due to the inherent smoothing effect of mean pooling, which emphasizes global consistency by averaging out local variations. In contrast, max pooling channels, which capture extreme activations, display higher variance as they remain more sensitive to spatial irregularities and outliers. Furthermore, across different experimental settings, a notable difference in variance was observed. In E1, where the test proteins were present in the training set, the kernel variance was relatively higher (except for the kernel associated with the mean pooling channel of the Metz dataset). This reflects the model’s ability to develop diverse and confident feature representations when processing familiar data. Conversely, in E3, where the test proteins were unseen during training, the kernel variance was relatively lower, indicating a more conservative and generalized response to out-of-distribution proteins. This suggests a shift in the spatial attention mechanism in E3, where the model relies more on broader global patterns rather than detailed local features, highlighting a potential generalization gap. These findings underscore the importance of enhancing kernel adaptability and feature diversity to improve the robustness of spatial attention layers, particularly in scenarios involving unseen data. By fostering greater adaptability and diversity in kernel activations, we can potentially mitigate the generalization gap and enhance the model’s performance on out-of-distribution samples.

**Figure 4.**
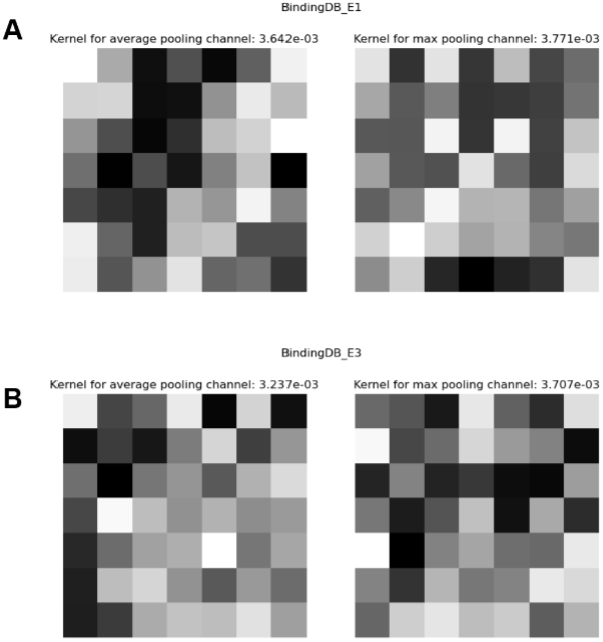
Kernel visualization of BindingDB dataset.

## 5 Discussion

One of the primary objectives of this study is to evaluate the significance of mHMG and iCBAM in drug-target interaction (DTI) prediction. Overall, our model demonstrated superior performance across all datasets. The ablation studies revealed that replacing mHMG with MG had a more substantial negative impact on the model’s performance compared to substituting iCBAM with spatial attention, particularly in regression tasks. This difference can be attributed to the greater length and diversity of protein sequences, which makes learning their patterns inherently more challenging. This observation also explains why LME proteins contributed more significantly to the model’s performance than LME molecules.

Furthermore, the overall performance of mHMG was slightly better than that of HMG, except in the KIBA dataset under E2. These results suggest that the incorporation of guiding knowledge by revising the HMG graph nodes enhances the model’s ability to extract meaningful information from molecular structures. A notable finding is the performance on the Metz dataset, where iCBAM outperformed spatial attention under E1 but performed worse under E3. In contrast, mHMG consistently outperformed both MG and HMG under E1, with the performance gap widening further under E2.

## 6 Conclusion

The investigation of DTIs is crucial for the development of new drugs. Although numerous deep learning-based approaches have been proposed to expedite this process, many methods suffer from two major limitations: the disregard for molecular motifs and the absence of spatial information regarding proteins. In this study, we address these challenges by employing mHMG for molecular encoding, which integrates motif features, and by utilizing an iCBAM to capture protein input features.

While earlier research suggested that the channel attention mechanism in CBAM could hinder performance, our findings indicate that modifying the sequential computation of spatial and channel attention into a parallel configuration within iCBAM enhances the model’s robustness and significantly improves its overall performance. To thoroughly evaluate the performance of our model, we designed four experimental settings using four datasets, addressing two types of tasks: classification task and regression task. Our comparative analysis with baseline models demonstrated that mHMG-iCBAM outperformed the baseline in 11 out of 16 experimental settings. Additionally, ablation experiments highlighted the importance of both mHMG and iCBAM in boosting the performance of our model. We believe our proposed model offers valuable insights for DTI prediction, presenting a more comprehensive and accurate approach to drug discovery.

## 7 Contribution

Zerui Yang initialized the project, implemented the model, and conducted the experiments. Linqi Song and Yudai Matsuda supervised the project. Linqi Song, Yudai Matsuda and Yinqiao Li critically reviewed the manuscript.

## 8 Acknowledgment

This work was supported in part by the Research Grants Council of the Hong Kong SAR under Grant GRF 11217823 and Collaborative Research Fund C1042-23GF, the National Natural Science Foundation of China under Grant 62371411, Technology and Innova tion Commission of Shenzhen Municipality under Grants JSGG20201102162000001, InnoHK initiative, the Government of the HKSAR, Laboratory for AI-Powered Financial Technologies.

