## Supplementary material for "mHMG-DTI: a drug-target interaction prediction framework combining modified Hierarchical Molecular Graphs and improved Convolutional Block Attention Module": SI

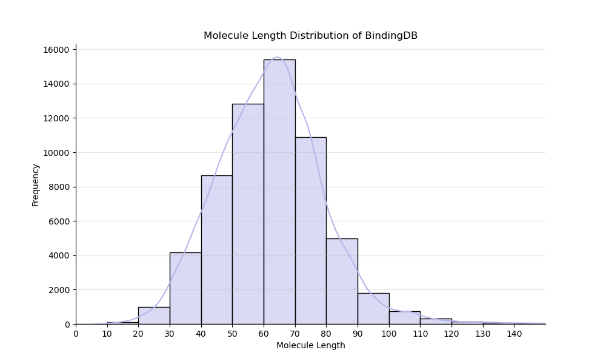

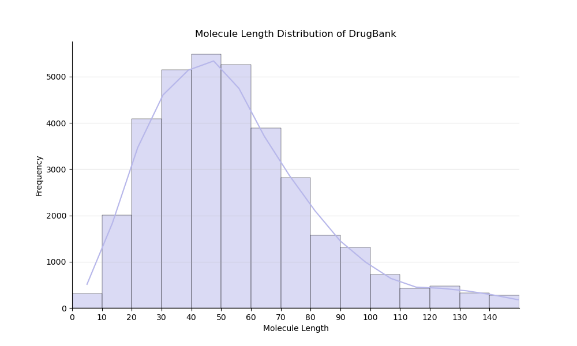

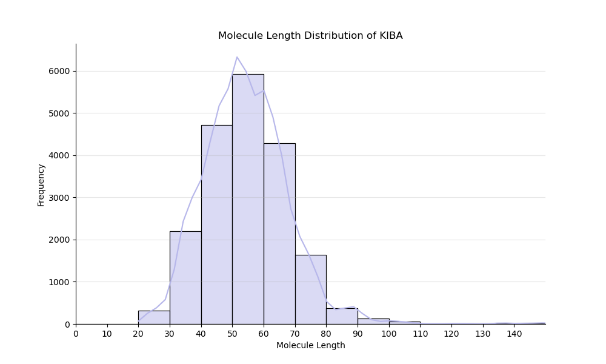

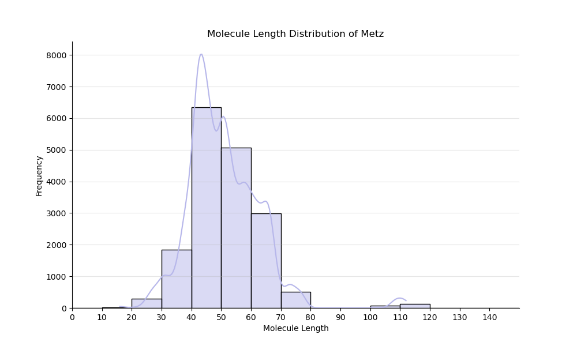

Figure S1: Molecule length of four datasets

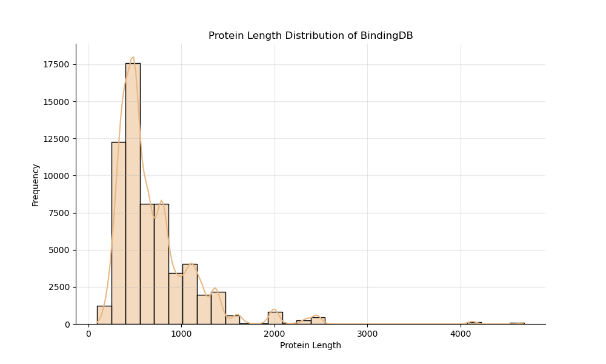

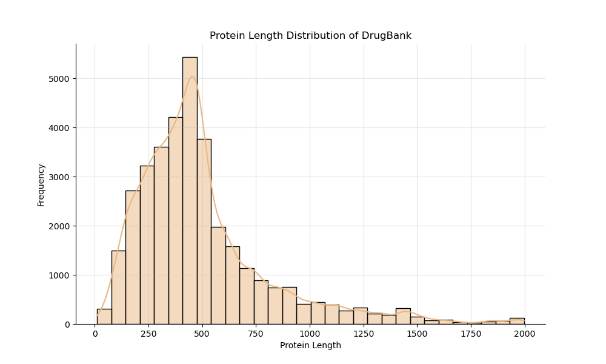

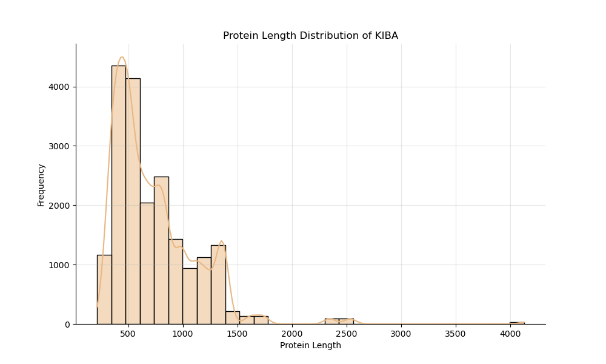

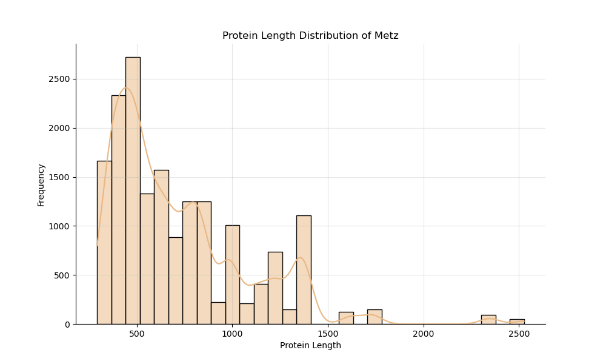

Figure S2: Protein length of four datasets

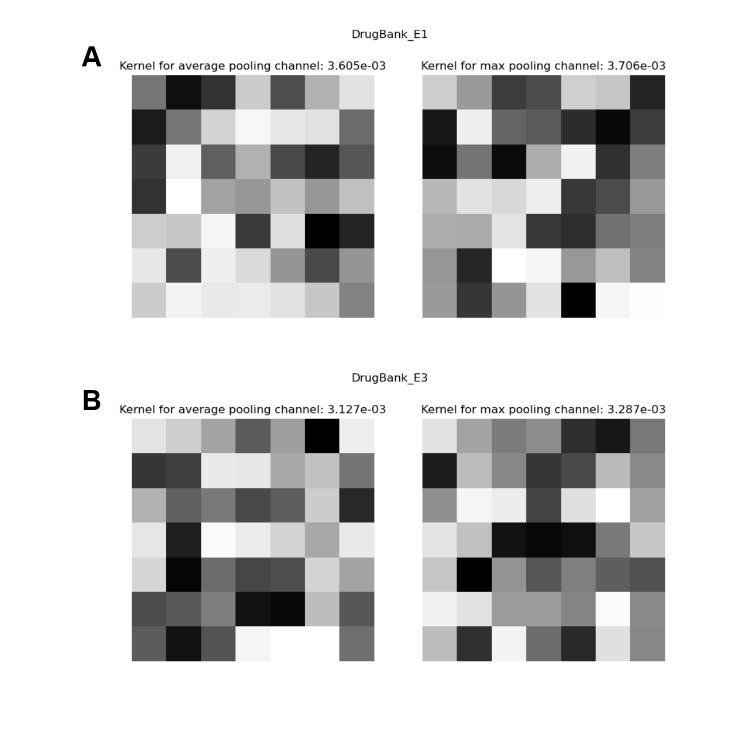

Figure S3: Kernel visualization of DrugBank dataset

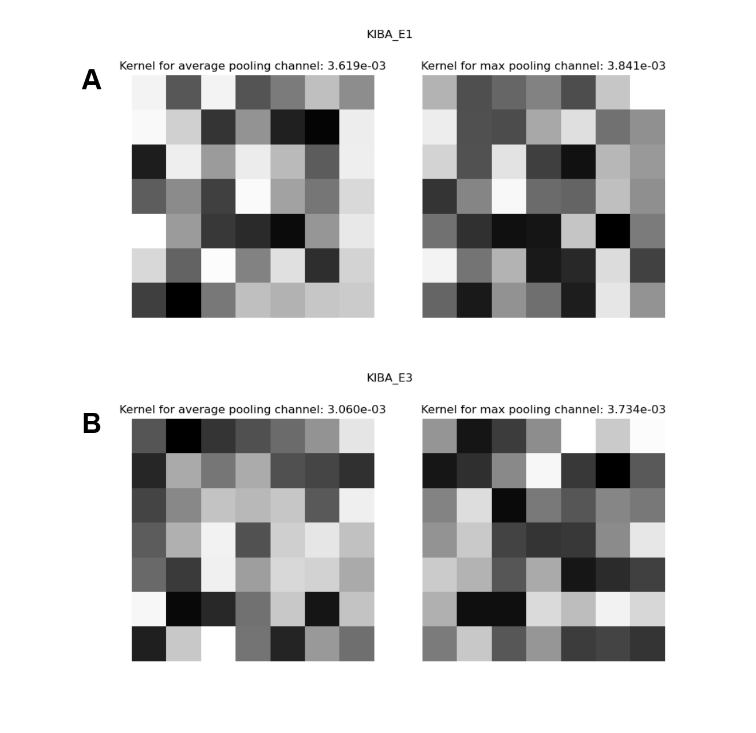

Figure S4: Kernel visualization of KIBA dataset

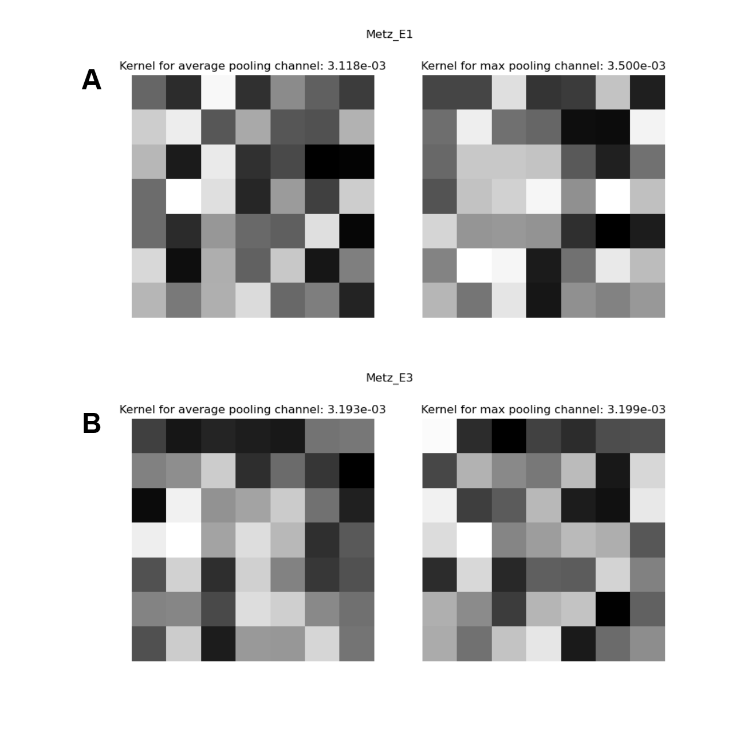

Figure S5: Kernel visualization of Metz dataset

| Models | ACC | PRE | Recall | MCC | F1 | ROC | PR |
| --- | --- | --- | --- | --- | --- | --- | --- |
| GraphDTA | 0.833 | 0.938 | 0.746 | 0.687 | 0.831 | 0.933 | 0.947 |
| DeepConv-DTI | 0.859 | 0.854 | 0.900 | 0.714 | 0.875 | 0.927 | 0.933 |
| HyperAttentionDTI | 0.921 | 0.913 | 0.947 | 0.841 | 0.930 | 0.975 | 0.980 |
| Perceiver CPI | 0.918 | 0.932 | 0.918 | 0.834 | 0.925 | 0.970 | 0.974 |
| HMG-DTI | 0.942 | 0.950 | 0.944 | 0.883 | 0.947 | 0.984 | 0.981 |

Table S1. Performance comparison on BindingDB dataset under E1

| Models | ACC | PRE | Recall | MCC | F1 | ROC | PR |
| --- | --- | --- | --- | --- | --- | --- | --- |
| GraphDTA | 0.858 | 0.938 | 0.800 | 0.728 | 0.864 | 0.952 | 0.960 |
| DeepConv-DTI | 0.902 | 0.920 | 0.902 | 0.801 | 0.911 | 0.963 | 0.967 |
| HyperAttentionDTI | 0.924 | 0.916 | 0.952 | 0.846 | 0.934 | 0.975 | 0.979 |
| Perceiver CPI | 0.935 | 0.934 | 0.951 | 0.869 | 0.943 | 0.980 | 0.984 |
| HMG-DTI | 0.947 | 0.947 | 0.959 | 0.892 | 0.953 | 0.985 | 0.987 |

Table S2. Performance comparison on BindingDB dataset under E2

| Models | ACC | PRE | Recall | MCC | F1 | ROC | PR |
| --- | --- | --- | --- | --- | --- | --- | --- |
| GraphDTA | 0.836 | 0.774 | 0.527 | 0.543 | 0.627 | 0.885 | 0.772 |
| DeepConv-DTI | 0.815 | 0.654 | 0.612 | 0.509 | 0.632 | 0.849 | 0.670 |
| HyperAttentionDTI | 0.896 | 0.787 | 0.826 | 0.736 | 0.806 | 0.925 | 0.834 |
| Perceiver CPI | 0.886 | 0.805 | 0.741 | 0.697 | 0.772 | 0.918 | 0.820 |
| HMG-DTI | 0.897 | 0.886 | 0.697 | 0.723 | 0.780 | 0.919 | 0.840 |

Table S3. Performance comparison on BindingDB dataset under E3

| Models | ACC | PRE | Recall | MCC | F1 | ROC | PR |
| --- | --- | --- | --- | --- | --- | --- | --- |
| GraphDTA | 0.552 | 0.879 | 0.387 | 0.286 | 0.538 | 0.714 | 0.830 |
| DeepConv-DTI | 0.630 | 0.764 | 0.651 | 0.227 | 0.703 | 0.667 | 0.786 |
| HyperAttentionDTI | 0.642 | 0.930 | 0.505 | 0.416 | 0.655 | 0.823 | 0.911 |
| Perceiver CPI | 0.723 | 0.823 | 0.745 | 0.403 | 0.784 | 0.766 | 0.844 |
| HMG-DTI | 0.682 | 0.724 | 0.767 | 0.326 | 0.745 | 0.721 | 0.751 |

Table S4. Performance comparison on BindingDB dataset under E4

| Models | ACC | PRE | Recall | MCC | F1 | ROC | PR |
| --- | --- | --- | --- | --- | --- | --- | --- |
| GraphDTA | 0.787 | 0.820 | 0.737 | 0.577 | 0.776 | 0.873 | 0.885 |
| DeepConv-DTI | 0.784 | 0.875 | 0.662 | 0.585 | 0.754 | 0.868 | 0.881 |
| HyperAttentionDTI | 0.867 | 0.844 | 0.898 | 0.734 | 0.871 | 0.840 | 0.863 |
| Perceiver CPI | 0.822 | 0.860 | 0.769 | 0.648 | 0.812 | 0.897 | 0.904 |
| HMG-DTI | 0.867 | 0.861 | 0.880 | 0.738 | 0.870 | 0.932 | 0.934 |

Table S5. Performance comparison on DrugBank dataset under E1

| Models | ACC | PRE | Recall | MCC | F1 | ROC | PR |
| --- | --- | --- | --- | --- | --- | --- | --- |
| GraphDTA | 0.689 | 0.771 | 0.518 | 0.395 | 0.619 | 0.744 | 0.771 |
| DeepConv-DTI | 0.707 | 0.805 | 0.529 | 0.435 | 0.638 | 0.777 | 0.789 |
| HyperAttentionDTI | 0.719 | 0.823 | 0.541 | 0.460 | 0.653 | 0.805 | 0.814 |
| Perceiver CPI | 0.740 | 0.733 | 0.737 | 0.480 | 0.734 | 0.823 | 0.831 |
| HMG-DTI | 0.778 | 0.783 | 0.755 | 0.556 | 0.769 | 0.853 | 0.859 |

Table S6. Performance comparison on DrugBank dataset under E2

| Models | ACC | PRE | Recall | MCC | F1 | ROC | PR |
| --- | --- | --- | --- | --- | --- | --- | --- |
| GraphDTA | 0.700 | 0.821 | 0.486 | 0.428 | 0.611 | 0.778 | 0.793 |
| DeepConv-DTI | 0.714 | 0.767 | 0.587 | 0.434 | 0.665 | 0.776 | 0.786 |
| HyperAttentionDTI | 0.742 | 0.776 | 0.655 | 0.486 | 0.710 | 0.808 | 0.816 |
| Perceiver CPI | 0.717 | 0.757 | 0.612 | 0.438 | 0.677 | 0.791 | 0.805 |
| HMG-DTI | 0.744 | 0.760 | 0.688 | 0.487 | 0.721 | 0.810 | 0.803 |

Table S7. Performance comparison on DrugBank dataset under E3

| Models | ACC | PRE | Recall | MCC | F1 | ROC | PR |
| --- | --- | --- | --- | --- | --- | --- | --- |
| GraphDTA | 0.576 | 0.585 | 0.457 | 0.151 | 0.513 | 0.594 | 0.611 |
| DeepConv-DTI | 0.615 | 0.794 | 0.289 | 0.283 | 0.424 | 0.665 | 0.691 |
| HyperAttentionDTI | 0.605 | 0.764 | 0.277 | 0.256 | 0.407 | 0.705 | 0.714 |
| Perceiver CPI | 0.644 | 0.703 | 0.472 | 0.299 | 0.565 | 0.685 | 0.702 |
| HMG-DTI | 0.646 | 0.731 | 0.438 | 0.312 | 0.548 | 0.722 | 0.712 |

Table S8. Performance comparison on DrugBank dataset under E4

| Models | MSE | CI |
| --- | --- | --- |
| GraphDTA | 0.695 | 0.685 |
| DeepConv-DTI | 0.486 | 0.715 |
| HyperAttentionDTI | 0.392 | 0.740 |
| Perceiver CPI | 0.395 | 0.753 |
| HMG-DTI | 0.361 | 0.768 |

Table S9. Performance comparison on KIBA dataset under E1

| Models | MSE | CI |
| --- | --- | --- |
| GraphDTA | 0.535 | 0.691 |
| DeepConv-DTI | 0.505 | 0.700 |
| HyperAttentionDTI | 0.506 | 0.692 |
| Perceiver CPI | 0.487 | 0.717 |
| HMG-DTI | 0.471 | 0.744 |

Table S10. Performance comparison on KIBA dataset under E2

| Models | MSE | CI |
| --- | --- | --- |
| GraphDTA | 0.458 | 0.607 |
| DeepConv-DTI | 0.418 | 0.625 |
| HyperAttentionDTI | 0.390 | 0.631 |
| Perceiver CPI | 0.396 | 0.614 |
| HMG-DTI | 0.410 | 0.582 |

Table S11. Performance comparison on KIBA dataset under E3

| Models | MSE | CI |
| --- | --- | --- |
| GraphDTA | 0.562 | 0.619 |
| DeepConv-DTI | 0.553 | 0.626 |
| HyperAttentionDTI | 0.555 | 0.608 |
| Perceiver CPI | 0.554 | 0.612 |
| HMG-DTI | 0.540 | 0.620 |

Table S12. Performance comparison on KIBA dataset under E4

| Models | MSE | CI |
| --- | --- | --- |
| GraphDTA | 0.425 | 0.713 |
| DeepConv-DTI | 0.344 | 0.753 |
| HyperAttentionDTI | 0.231 | 0.811 |
| Perceiver CPI | 0.222 | 0.815 |
| HMG-DTI | 0.186 | 0.841 |

Table S13. Performance comparison on Metz dataset under E1

| Models | MSE | CI |
| --- | --- | --- |
| GraphDTA | 0.404 | 0.691 |
| DeepConv-DTI | 0.307 | 0.688 |
| HyperAttentionDTI | 0.244 | 0.748 |
| Perceiver CPI | 0.230 | 0.760 |
| HMG-DTI | 0.252 | 0.779 |

Table S14. Performance comparison on Metz dataset under E2

| Models | MSE | CI |
| --- | --- | --- |
| GraphDTA | 0.577 | 0.669 |
| DeepConv-DTI | 0.441 | 0.708 |
| HyperAttentionDTI | 0.348 | 0.752 |
| Perceiver CPI | 0.399 | 0.694 |
| HMG-DTI | 0.452 | 0.678 |

Table S15. Performance comparison on Metz dataset under E3

| Models | MSE | CI |
| --- | --- | --- |
| GraphDTA | 0.704 | 0.634 |
| DeepConv-DTI | 0.660 | 0.616 |
| HyperAttentionDTI | 0.570 | 0.669 |
| Perceiver CPI | 0.555 | 0.616 |
| HMG-DTI | 0.541 | 0.689 |

Table S16. Performance comparison on Metz dataset under E4
